# Interferon-β Coordinates Epithelial Immune Networks and Fibrotic Responses During *Chlamydia muridarum* Infection

**DOI:** 10.64898/2026.03.23.713583

**Authors:** Ramesh Kumar, Iliana C. Cordova-Mendez, Derrick Burgess, Brahim Qadadri, Arkaprabha Banerjee, Wilbert A. Derbigny

**Affiliations:** Department of Microbiology, Marian University-Wood College of Osteopathic Medicine Indianapolis, Indiana 46222; Department of Pediatrics; Dept. of Microbiology; Dept. of Infectious Diseases; Indiana University School of Medicine Indianapolis, Indiana 46202

**Keywords:** Oviduct epithelial cells, *Chlamydia*, IFNβ-deficiency, cytokines, epithelial, mucosa, pathway-focused qPCR assays, inflammation, fibrosis, innate immunity

## Abstract

*Chlamydia trachomatis* infection is the most common bacterial sexually transmitted infection worldwide and a leading cause of inflammatory reproductive tract disease and infertility in women. Much of the tissue damage associated with genital chlamydial infection arises from host inflammatory responses rather than direct bacterial cytotoxicity. Epithelial cells lining the female reproductive tract represent the primary host cells infected during chlamydial infection and play key roles in initiating innate immune responses. Among the cytokines produced by infected epithelial cells, type-I interferons have emerged as important regulators of host defense and inflammatory signaling; however, the specific contribution of interferon-β (IFN-β) to epithelial transcriptional responses during chlamydial infection remains incompletely defined.

In the present study, we investigated the role of IFN-β in coordinating epithelial immune signaling networks during infection with Chlamydia muridarum. Using wild-type murine oviduct epithelial cells (OE-WT) and IFN-β-deficient epithelial cells (OE-IFNβ-KO), we performed pathway-focused RT² Profiler PCR array analyses examining transcriptional responses across four biological pathways: (1) innate and adaptive immune responses, (2) type-I interferon signaling, (3) inflammatory and autoimmune responses, and (4) fibrosis-associated pathways. Infection of OE-WT cells resulted in coordinated induction of cytokines, chemokines, and interferon-stimulated genes associated with antimicrobial defense and immune cell recruitment. In contrast, IFN-β deficiency resulted in widespread dysregulation of these transcriptional programs, including reduced induction of interferon-responsive chemokines such as CCL5 and CXCL10, altered inflammatory cytokine expression, and transcriptional signatures consistent with enhanced tissue remodeling responses.

Notably, IFN-β deficiency resulted in increased TNF expression accompanied by reduced IL-6 induction, suggesting disruption of balanced inflammatory signaling networks. Pathway analyses further revealed dysregulated expression of fibrosis-associated genes including *Serpine1, Ctgf,* and *Eng* in IFN-β-deficient epithelial cells, indicating potential mechanisms linking interferon signaling to tissue remodeling during infection. Collectively, these findings identify IFN-β as a central regulator of epithelial immune networks during chlamydial infection and suggest that disruption of IFN-β signaling may promote inflammatory and fibrotic pathology within the female reproductive tract.

**Author Summary:** Sexually transmitted infections caused by *Chlamydia trachomatis* are a major cause of infertility worldwide. Although antibiotic treatment can eliminate the bacteria, damage to the reproductive tract often results from the body’s own immune response to infection. The epithelial cells lining the reproductive tract are the first cells infected and play an important role in initiating immune responses. In this study, we investigated how a specific immune signaling molecule, interferon-β (IFN-β), regulates the gene expression programs activated in epithelial cells during chlamydial infection. Using pathway-focused gene expression arrays, we found that IFN-β coordinates multiple immune pathways, including interferon signaling, inflammatory cytokine networks, and genes associated with tissue remodeling. When IFN-β was absent, many of these pathways became dysregulated, resulting in altered inflammatory signaling and gene expression patterns linked to fibrosis. These findings suggest that IFN-β functions as a key regulator that helps balance protective immune responses with inflammatory processes that can damage reproductive tissues during infection.

## Introduction

*Chlamydia trachomatis* is the most frequently reported bacterial sexually transmitted infection worldwide and remains a major cause of inflammatory reproductive tract disease and infertility. The World Health Organization estimates that more than 120 million new infections occur globally each year, with a large proportion of infections occurring in women of reproductive age (1). Although antibiotic treatment is effective at eliminating the pathogen, many infections remain asymptomatic, untreated or recurrent infections can lead to serious complications including pelvic inflammatory disease, ectopic pregnancy, and tubal factor infertility (2).

A central feature of chlamydial pathogenesis is the complex relationship between protective host immunity and inflammatory responses that drive tissue damage. While host immune responses are required for controlling infection, excessive or dysregulated inflammation is believed to be the primary driver of tissue damage within the reproductive tract (3). Chronic inflammation within the fallopian tubes can lead to fibrosis and scarring that ultimately compromises reproductive function. Understanding the molecular mechanisms that regulate this balance between antimicrobial defense and immunopathology remains a critical challenge in the field.

Epithelial cells lining the female reproductive tract serve as the primary cellular targets of chlamydial infection and represent the first line of host defense against invading pathogens. Infected epithelial cells produce numerous cytokines and chemokines that initiate innate immune responses, induce an inflammatory response within infected tissues, and recruit leukocytes to sites of infection (4, 5). These epithelial-derived mediators play critical roles in shaping subsequent immune responses, including activation of innate immune pathways and modulation of adaptive immunity. Because epithelial cells represent both the primary replicative niche for *Chlamydia* and the earliest source of inflammatory signals, the transcriptional programs activated in infected epithelial cells are likely to be critical determinants of disease outcome.

Among the cytokines produced during infection, type-I interferons have emerged as important regulators of host responses to intracellular pathogens. Type-I interferons, including interferon-α (IFN-α) and interferon-β (IFN-β), activate transcriptional programs that influence antiviral defenses, inflammatory responses, and adaptive immunity (6). However, the role of type-I interferons during bacterial infections is complex, context-dependent, and often paradoxical, with studies demonstrating both protective and pathogenic effects depending on the pathogen and tissue environment.

Previous studies from our laboratory and others have demonstrated that *Chlamydia muridarum* infection induces IFN-β production in epithelial cells through activation of innate immune sensing pathways including Toll-like receptor 3 (TLR3) signaling (7-9). Interestingly, genetic deficiency of TLR3 results in increased bacterial burden and exacerbated genital tract pathology in experimental models of chlamydial infection, suggesting that TLR3-dependent pathways contribute to protective host responses during infection (7, 8, 10, 11). TLR3 deficiency reduces chlamydial induction of IFN-β in OE cells by approximately 60–70%, indicating that additional signaling pathways contribute to IFN-β production during infection. However, in experiments in which TLR3-deficient OE cells were supplemented with purified exogenous IFN-β prior to infection, we observed significant reductions in chlamydial replication and the size of the chlamydial inclusion in these cells (12). These findings suggest that IFN-β may have some role in limiting chlamydial replication *in vitro*. Understanding how epithelial cell type-1 interferon signaling coordinates inflammatory and tissue-remodeling pathways is critical for defining the molecular mechanisms linking chlamydial infection to reproductive tract pathology.

Despite a growing recognition of the importance of interferon signaling during chlamydial infection, the downstream transcriptional networks regulated by IFN-β in epithelial cells remain poorly understood. However, it remains unclear how IFN-β signaling integrates with inflammatory cytokine pathways and tissue remodeling programs that ultimately influence disease outcomes during infection. In the present study, we investigated the role of IFN-β in coordinating epithelial transcriptional responses during *Chlamydia muridarum* infection. Using pathway-focused RT² Profiler PCR arrays, we examined transcriptional responses across multiple immune pathways in wild-type murine oviduct epithelial (OE) cells and IFNβ-deficient OE cells following infection. Our analysis demonstrates that IFN-β functions as a central regulatory node coordinating interferon signaling, inflammatory cytokine networks, and fibrosis-associated pathways.

These findings provide new insight into how interferon signaling regulates epithelial immune responses and suggest mechanisms linking dysregulated interferon pathways to reproductive tract pathology during chlamydial infection.

## Materials and Methods

### Cell lines and culture conditions

Murine oviduct epithelial (OE) cell-lines were derived from wild-type C57BL6 and interferon-β-deficient mice generated on a C57BL/6 background (13) as previously described for other OE cell lines generated in our lab (5, 9, 14). Once isolated and clonally expanded, wild-type (OE-WT) and interferon-β-deficient (OE-IFNβ-KO) OE cells were maintained and cultured in epithelial cell media comprised of a 1:1 ratio DMEM:F12K (Sigma-Aldrich), supplemented with 10% characterized FBS (Thermo-Fisher, Pittsburgh, PA), 2 mM L-alanyl-L-glutamine (GlutaMAX I; Life Technologies/Invitrogen, Carlsbad, CA), 5 mg bovine insulin/ml, and 12.5 ng/ml recombinant human fibroblast growth factor-7 (keratinocyte growth factor; Sigma-Aldrich). OE cells are cultured and maintained at 37°C in a humidified incubator with 5% CO₂ for all experiments.

### Chlamydia infection

*Mycoplasma*-free *Chlamydia muridarum* (strain Nigg) propagated in McCoy cells, as described previously in (15) was used for all infections. OE cells were infected at a multiplicity of infection (MOI) of 10 as described previously (5, 9, 14). Infections were performed by inoculating epithelial cell monolayers with bacterial suspensions followed by centrifugation-assisted infection to facilitate bacterial entry. After centrifugation, cultures were incubated under standard culture conditions for the indicated time points before cells were harvested and total cell RNA was isolated.

### RNA isolation and cDNA synthesis

Total RNA was isolated using the Qiagen RNeasy Plus Mini Kit (Qiagen, Valencia, CA) according to the manufacturer’s instructions. During purification, all RNA samples were treated with column-based chromatography (Qiagen) to remove genomic-DNA contamination. RNA concentrations and purity were determined using spectrophotometric analysis. Reverse transcription reactions were performed using the Qiagen RT² First Strand Kit with 500 ng total RNA per reaction according to the manufacturer’s protocol.

### RT² Profiler PCR array analysis and Quantitative real-time PCR

Gene expression was analyzed using Qiagen RT² Profiler PCR Arrays designed to examine transcriptional responses across specific biological pathways. The following pathway-focused arrays were used: <u>Innate and Adaptive Immune Responses</u> (PAMM-052Z), <u>Type-I Interferon Responses</u> (PAMM-016Z), <u>Fibrosis Pathways</u> (PAMM-120Z), and <u>Inflammatory Response and Autoimmunity</u> (PAMM-077Z). Each array contains gene targets associated with key components of the corresponding biological pathway. Quantitative real-time PCR was conducted with the diluted cDNA and gene-specific primers according to the protocol outlined in the iTaq SYBR green Supermix with ROX kit (Bio-Rad, Hercules,CA) and as described previously (9). Dissociation curves were recorded after each run to ensure primer specificity.

### Quantitative PCR instrumentation

PCR reactions were performed using an Applied Biosystems QuantStudio 3 Real-Time PCR System (Applied Biosystems, Norwalk, CT), according to manufacturer recommendations.

### Data normalization and analysis

Gene expression levels in the Qiagen RT² Profiler PCR Arrays were normalized using the following housekeeping genes included on the arrays*: Actb, B2m, Gapdh, Gusb*, and *Hsp90ab1*. Relative gene expression was calculated using the ΔΔCt method. Fold regulation values were determined relative to mock-infected controls. Genes exhibiting ≥2-fold change with *p < 0.05* were considered significantly regulated.

### Statistical Analysis

Numerical data are presented as mean ± SD. <u>All</u> experiments were performed in triplicate and repeated at least twice for the RT² Profiler PCR Arrays, and three times for standard qPCR. The data were analyzed using appropriate statistical tests as indicated in the figure legends. Data were tested for normality prior to parametric analysis. Comparisons between groups were performed using Student’s *t* test or analysis of variance (ANOVA) in GraphPad Prism where appropriate. A *p*-value of <0.05 was considered statistically significant.

## RESULTS

### IFN-β regulates innate and adaptive immune transcriptional responses during *Chlamydia muridarum* infection

To investigate the role of IFN-β in epithelial cell immune responses during chlamydial infection, we first analyzed transcriptional changes in genes associated with innate and adaptive immune signaling using the RT² Profiler PCR array for <u>innate and adaptive immune responses</u> (PAMM-052Z). OE-WT and OE-IFNβ-KO cells were infected with *C. muridarum* (MOI = 10), and gene expression profiles were examined at 18 and 32 hours post-infection, corresponding to middle and late stages of the chlamydial developmental cycle.

In OE-WT cells, infection induced robust transcriptional activation of multiple immune mediators, including cytokines and chemokines involved in leukocyte recruitment and inflammatory signaling (**Figure 1**). Several genes exhibited strong induction at both time points, including *Ccl5, Cxcl10, Il6, and Ifnγ,* indicating activation of epithelial inflammatory signaling networks in response to infection.

**Figure 1.**
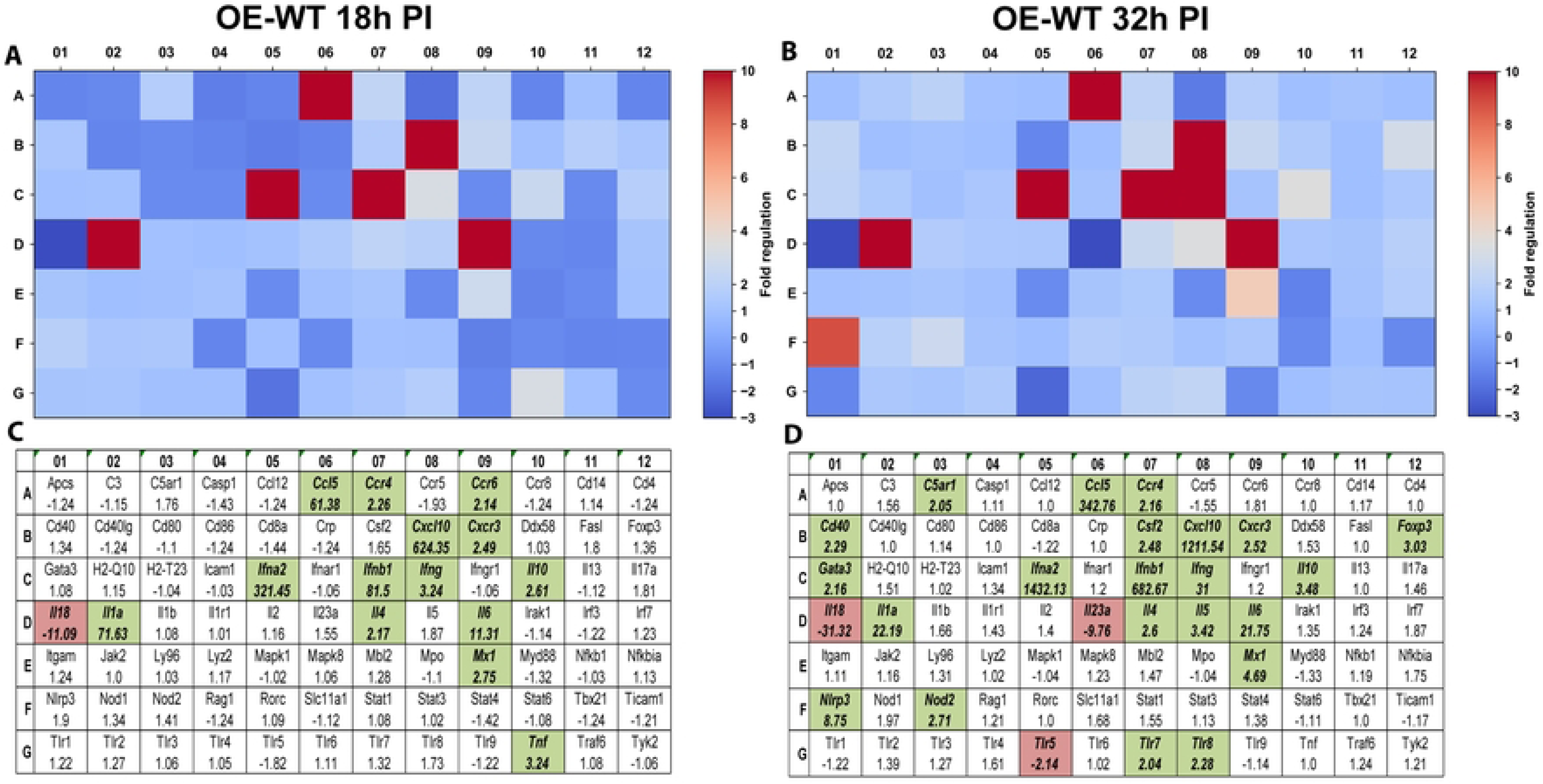
RT^2^ Profiler pathway-focused qPCR analyses of various genes associated with the innate and adaptive immune response. Gene-expression heat-maps and gene charts of wild-type oviduct epithelial cells (OE-WT) that were infected with *C. muridarum* at an MOI=10 for either 18hrs (A, C) or 32hrs (B, D) post-infection (PI). The data shown are representative and show fold changes in gene expression levels relative to mock-infected controls. The colors green and red indicate at least a 2-fold increase or decrease in gene expression, respectively.

In contrast, IFN-β deficiency significantly altered the transcriptional response to infection (**Figure 2**). Several genes strongly induced in OE-WT cells exhibited markedly reduced induction in OE-IFNβ-KO cells, including the interferon-responsive chemokines *Ccl5* and *Cxcl10*. These findings indicate that IFN-β signaling contributes to the induction of epithelial chemokine networks that coordinate immune cell recruitment during infection. Interestingly, not all transcriptional responses were reduced in the absence of IFN-β. Some inflammatory mediators, including *Tnf, Il23a, and Il1β,* showed increased expression in IFNβ-deficient cells compared with wild-type controls, suggesting that IFN-β may also help restrain specific inflammatory pathways during infection. Collectively, these findings indicate that IFN-β plays an important role in coordinating immune signaling networks in epithelial cells during chlamydial infection of murine OE cells.

**Figure 2.**
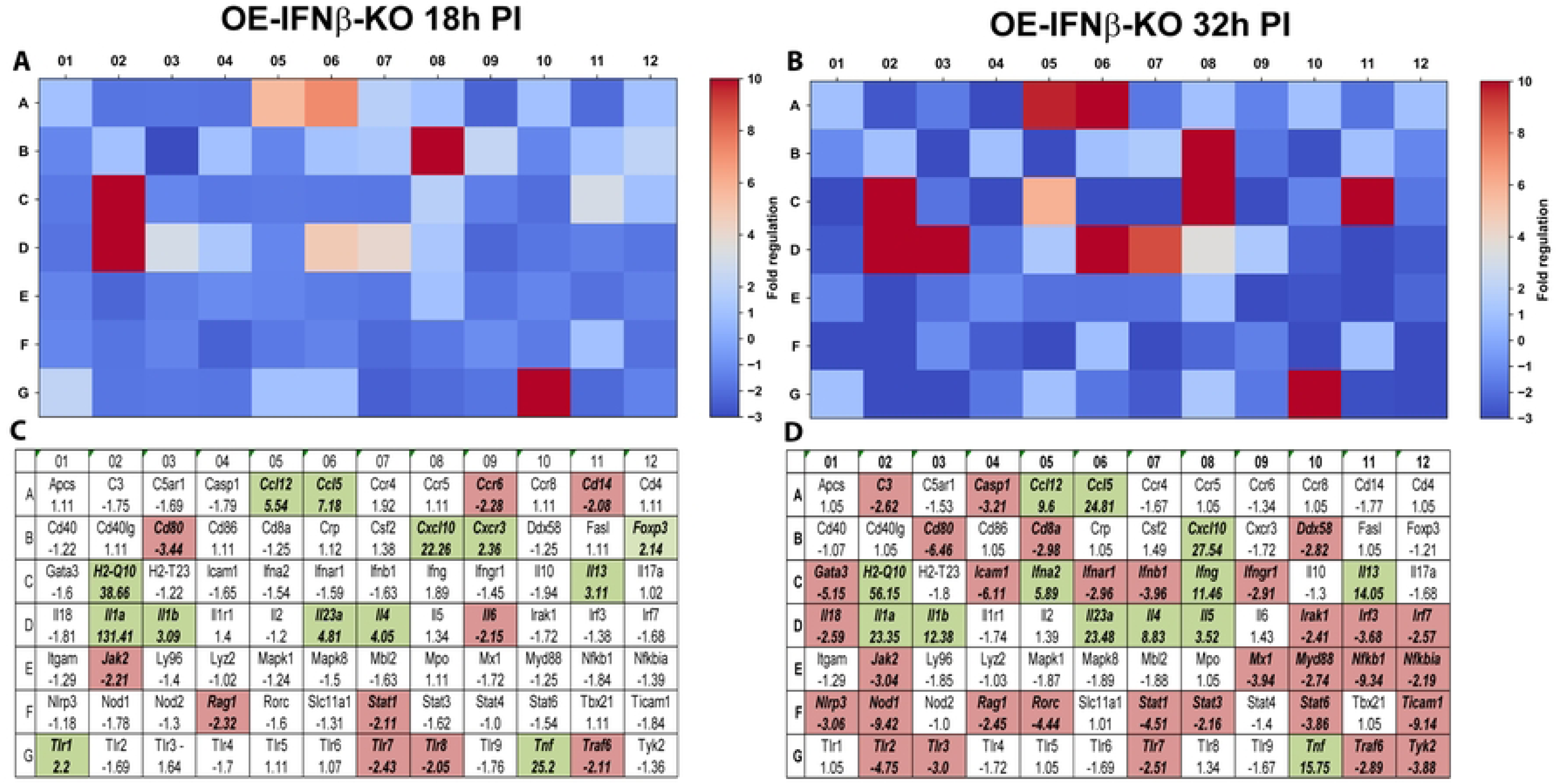
RT^2^ Profiler pathway-focused qPCR analyses of various genes associated with the innate and adaptive immune response during IFNβ deficiency. Gene-expression heat-maps and gene charts of IFNβ-knockout epithelial cells (OE-IFNβ-KO) that were infected with *C. muridarum* at an MOI=10 for either 18hrs (A, C) or 32hrs (B, D) post-infection (PI). The data shown are representative and show fold changes in gene expression levels relative to mock-infected controls. The colors green and red in the gene charts (C and D) indicate at least 2-fold increase or decrease, respectively, in gene expression levels.

### IFN-β amplifies type-I interferon transcriptional programs during infection

We next examined transcriptional responses within the type-I interferon signaling pathway using the RT² Profiler PCR array for <u>type-I interferon responses</u> (PAMM-016Z). Gene expression profiles were analyzed at 12 and 24 hours post-infection, corresponding to early-middle and late stages of the chlamydial developmental cycle. In OE-WT cells, infection induced strong transcriptional activation of numerous interferon-stimulated genes (ISGs), including *Isg15, Oas1, Ifit1, Ifit3, Mx1,* and *Stat1* (**Figure 3**). Many of these genes are well-characterized components of interferon-mediated antimicrobial defense pathways and are known to be induced downstream of type-I interferon signaling (16).

**Figure 3.**
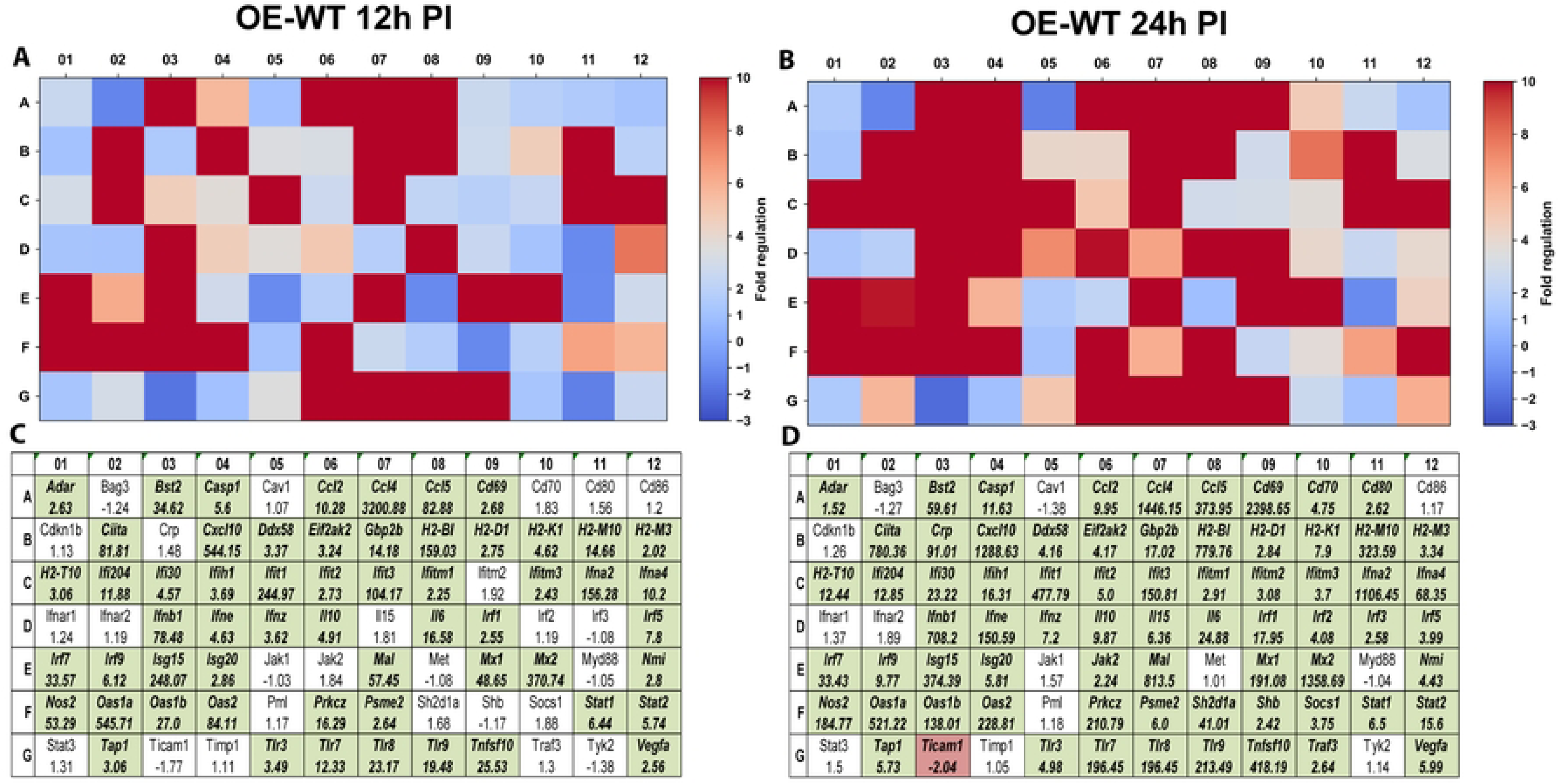
RT^2^ Profiler pathway-focused qPCR analyses of various genes associated with the type-1 IFN. Gene-expression heat-maps and gene charts of wild-type oviduct epithelial cells (OE-WT) that were infected with *C. muridarum* at an MOI=10 for either 12hrs (A, C) or 24hrs (B, D) post-infection (PI). The data shown are representative and show fold changes in gene expression levels relative to mock-infected controls. The colors green and red in the gene charts (C and D) indicate at least a 2-fold increase or decrease, respectively, in gene expression levels.

In contrast, OE-IFNβ-KO cells exhibited markedly attenuated induction of many interferon-stimulated genes (**Figure 4**). Several genes that were strongly induced in OE-WT cells exhibited minimal or absent induction in IFNβ-deficient cells, indicating that IFN-β functions as a key amplifier of interferon signaling during infection. Notably, the magnitude of transcriptional attenuation varied across different genes. Some interferon-responsive genes exhibited near-complete loss of induction, whereas others displayed partial reductions in expression, suggesting that additional signaling pathways may contribute to interferon-stimulated gene activation during infection. These findings are consistent with previous observations demonstrating that TLR3 signaling contributes substantially, but not exclusively, to IFN-β induction during chlamydial infection (9). Together, these data indicate that IFN-β functions within a broader network of innate immune signaling pathways that regulate interferon-stimulated gene expression in epithelial cells.

**Figure 4.**
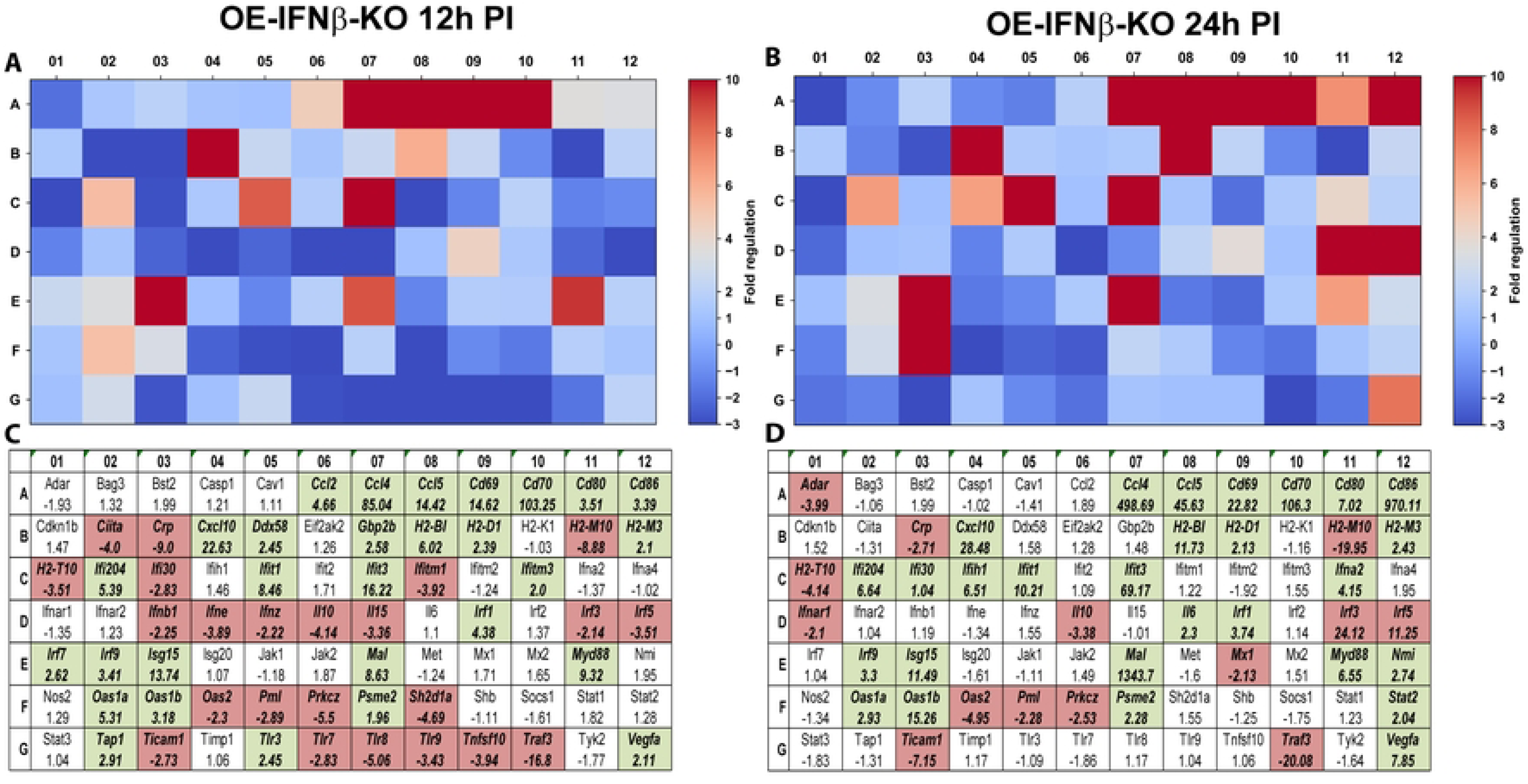
RT^2^ Profiler pathway-focused qPCR analyses of various genes associated with type-1 IFN during IFNβ deficiency. Gene-expression heat-maps and gene charts of IFNβ-knockout epithelial cells (OE-IFNβ-KO) that were infected with *C. muridarum* at an MOI=10 for either 12hrs (A, C) or 24hrs (B, D) post-infection (PI). The data shown are representative and show fold changes in gene expression levels relative to mock-infected controls. The colors green and red in the gene charts (C and D) indicate at least a 2-fold increase or decrease, respectively, in gene expression levels.

### IFN-β modulates epithelial transcriptional programs associated with tissue remodeling and fibrosis

Because chronic inflammation during genital tract chlamydial infection is strongly associated with fibrosis and scarring within the reproductive tract, we next examined transcriptional changes in genes associated with fibrosis and extracellular matrix remodeling using the RT² Profiler PCR array for <u>fibrosis pathways</u> (PAMM-120Z). In OE-WT cells, infection induced transcriptional changes in several genes associated with extracellular matrix remodeling, turnover, and tissue repair, including *Mmp9, Col3a1, Timp1,* and *Serpine1* (**Figure 5**). These genes play important roles in regulating extracellular matrix dynamics during inflammatory responses.(17-20).

**Figure 5.**
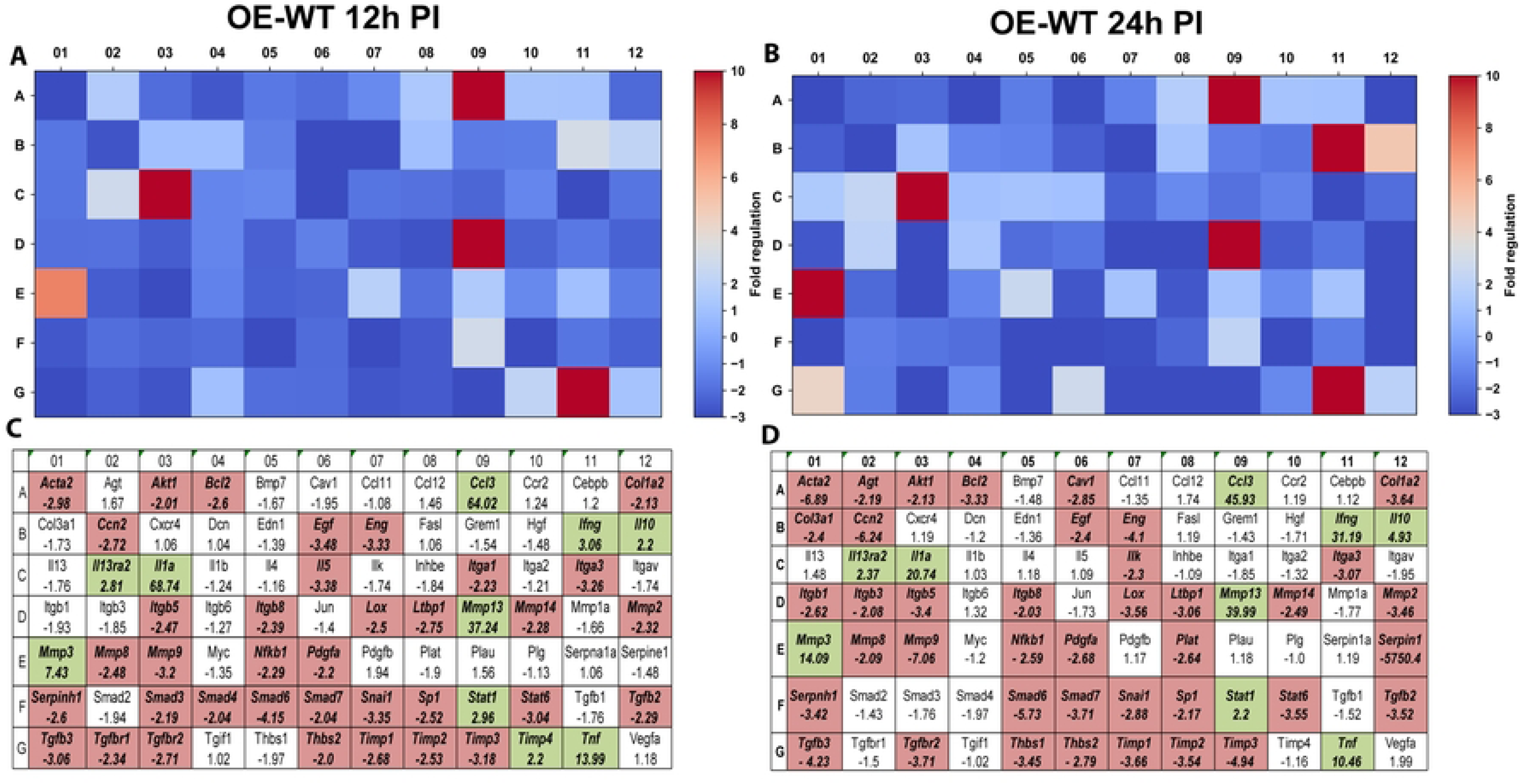
RT^2^ Profiler pathway-focused qPCR analyses of various genes associated with fibrosis. Gene-expression heat-maps and gene charts of wild-type oviduct epithelial cells (OE-WT) that were infected with *C. muridarum* at an MOI=10 for either 12hrs (A, C) or 24hrs (B, D) post-infection (PI). The data shown are representative and show fold changes in gene expression levels relative to mock-infected controls. The colors green and red in the gene charts (C and D) indicate at least a 2-fold increase or decrease, respectively, in gene expression levels.

In contrast, IFN-β deficiency resulted in substantial alterations in fibrosis-associated gene expression (**Figure 6**). Several genes associated with pro-fibrotic signaling, including *Ctgf (Ccn2), Eng,* and *Tnf*, exhibited increased expression in OE-IFNβ-KO cells relative to wild-type controls. These genes have been implicated in fibroblast activation and extracellular matrix deposition in multiple inflammatory disease models (21-24). Additionally, several matrix-metalloproteinases (MMPs) exhibited increased expression in the absence of IFN-β. MMPs are a family of zinc-dependent enzymes that degrade and remodel extracellular matrix (ECM) proteins. Dysregulation of MMP transcription is known to contribute to chronic wounds, tumor metastasis, and fibrosis associated with various inflammatory diseases (25-27).

**Figure 6.**
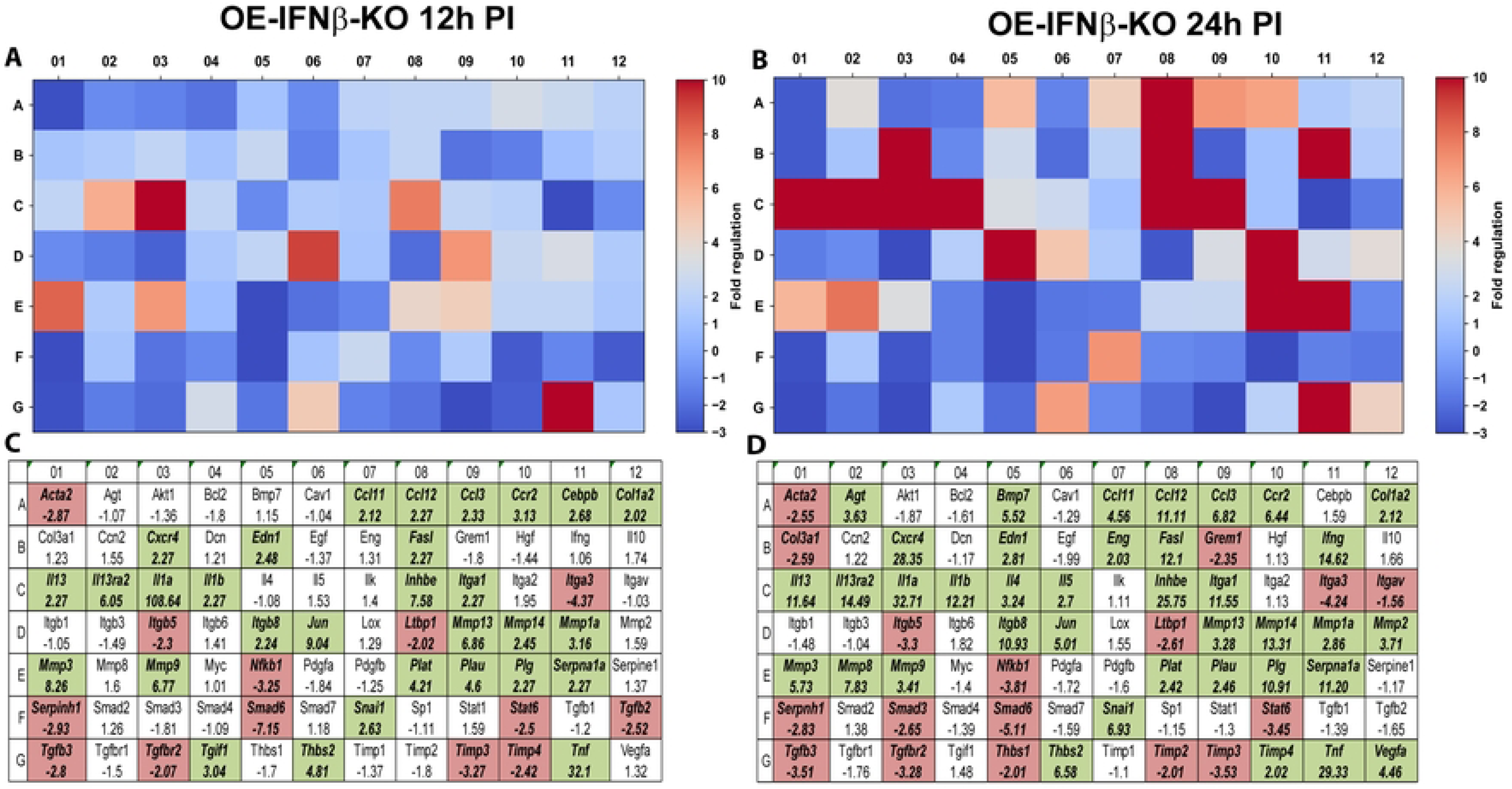
RT^2^ Profiler pathway-focused qPCR analyses of various genes associated with fibrosis during IFNβ deficiency. Gene-expression heat-maps and gene charts of IFNβ-knockout epithelial cells (OE-IFNβ-KO) that were infected with *C. muridarum* at an MOI=10 for either 12hrs (A, C) or 24hrs (B, D) post-infection (PI). The data shown are representative and show fold changes in gene expression levels relative to mock-infected controls. The colors green and red in the gene charts (C and D) indicate at least a 2-fold increase or decrease, respectively, in gene expression levels.

Interestingly, other genes exhibited decreased expression in IFNβ-deficient cells, indicating that IFN-β regulates multiple aspects of the tissue remodeling response. For example, the regulation of *Serpine1*, which encodes plasminogen activator inhibitor-1, differed markedly between wild-type and IFNβ-deficient cells. In OE-WT cells, *Serpine1* expression was strongly reduced at later time points during infection, approaching the assay’s detection threshold. In contrast, *Serpine1* expression remained detectable in IFNβ-deficient cells. Because PAI-1 is a key regulator of fibrinolysis and extracellular matrix turnover, altered regulation of this gene may influence tissue remodeling responses during infection.

Collectively, these findings indicate that IFN-β signaling influences multiple transcriptional pathways associated with extracellular matrix remodeling and fibrotic signaling during chlamydial infection. Dysregulation of these pathways due to the reduction or absence of IFN-β may contribute to enhanced tissue pathology observed in the experimental models of chlamydial infection.

### IFN-β coordinates epithelial inflammatory signaling networks

To further investigate inflammatory transcriptional responses during infection, we examined gene expression changes within pathways associated with <u>inflammatory and autoimmune responses</u> using the RT² Profiler PCR array (PAMM-077Z). In OE-WT cells, infection induced strong transcriptional activation of numerous inflammatory mediators, including *Ccl5, Ccl2, Cxcl10, Il1β,* and *Tnf* (**Figure 7**). These cytokines and chemokines play key roles in orchestrating leukocyte recruitment and inflammatory responses during infection (28-32).

**Figure 7.**
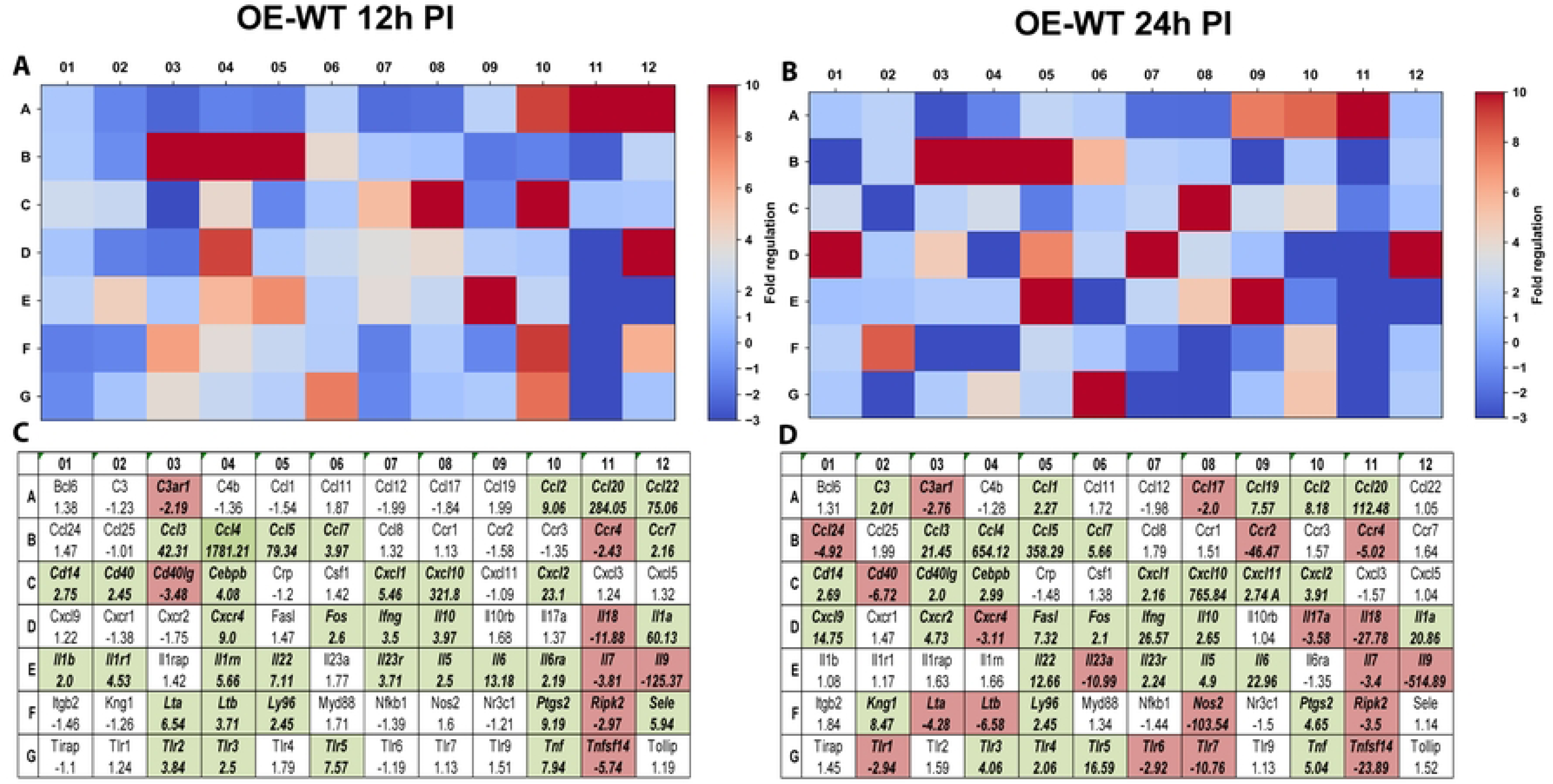
RT^2^ Profiler pathway-focused qPCR analyses of various genes associated with the inflammatory response & autoimmunity. Gene-expression heat-maps and gene charts of wild-type oviduct epithelial cells (OE-WT) that were infected with *C. muridarum* at an MOI=10 for either 12hrs (A, C) or 24hrs (B, D) post-infection (PI). The data shown are representative and show fold changes in gene expression levels relative to mock-infected controls. The colors green and red in the gene charts (C and D) indicate at least a 2-fold increase or decrease, respectively, in gene expression levels.

In contrast, IFN-β deficiency resulted in complex alterations in inflammatory gene expression (**Figure 8**). Several interferon-associated chemokines, including *Ccl5* and *Cxcl10,* exhibited reduced induction in OE-IFNβ-KO cells, consistent with their regulation by interferon signaling pathways. However, other inflammatory mediators exhibited divergent expression levels in IFN-β-deficient cells. For example, *Tnf* was substantially elevated in the OE-IFNβ-KO cells relative to wild-type controls. In contrast, *Il6* expression was reduced in IFNβ-deficient cells, suggesting that IFN-β may differentially regulate specific inflammatory pathways during infection. Interestingly, the transcription of chronic inflammatory markers such as Il23a and Il23r was significantly increased during interferon-β deficiency. Dysregulation of IL-23A and IL-23R transcription is a significant factor in the pathogenesis of chronic inflammation and autoimmunity (33). Together, these findings indicate that IFN-β coordinates complex inflammatory signaling networks during chlamydial infection and may help balance protective immune responses with inflammatory pathways that contribute to tissue pathology, chronic inflammation, and autoimmunity.

**Figure 8.**
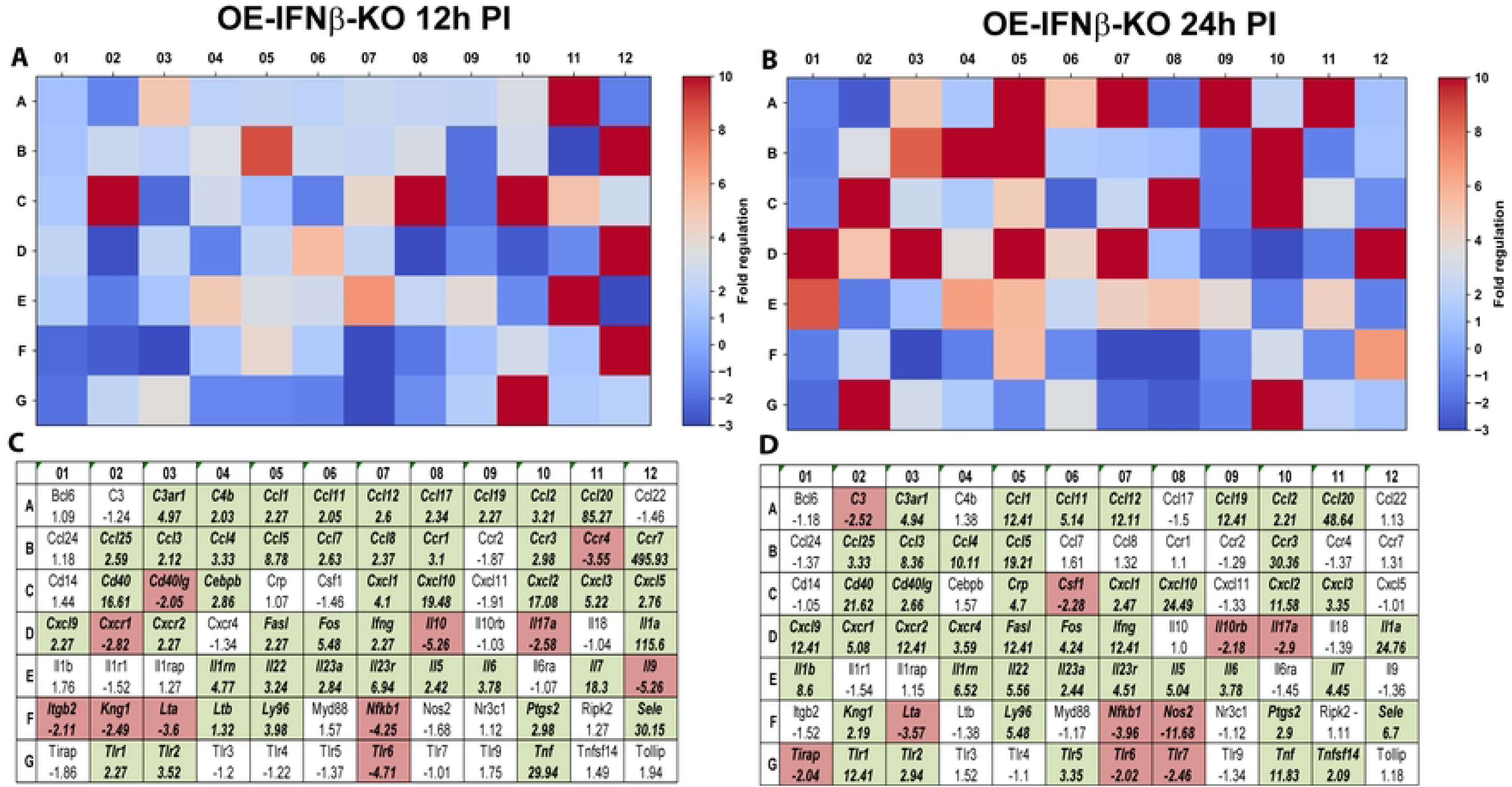
RT^2^ Profiler pathway-focused qPCR analyses of various genes associated with the inflammatory response and autoimmunity during IFNβ deficiency. Gene-expression heat-maps and gene charts of IFNβ-knockout epithelial cells (OE-IFNβ-KO) that were infected with *C. muridarum* at an MOI=10 for either 12hrs (A, C) or 24hrs (B, D) post-infection (PI). The data shown are representative and show fold changes in gene expression levels relative to mock-infected controls. The colors green and red in the gene charts (C and D) indicate at least a 2-fold increase or decrease, respectively, in gene expression levels.

### Validation of selected genes by independent qPCR

To validate the transcriptional changes identified in the pathway-focused PCR arrays, we performed independent quantitative PCR analyses on selected genes representing key inflammatory and immune signaling pathways. Independent qPCR analyses confirmed the differential expression patterns observed in the array datasets, including altered regulation of *Ccl5, Cxcl10, Cxcl16, IFN-γ, Mmp9, Tnf, Il4,* and *Il6* in IFN-β-deficient cells relative to wild-type controls (**Figures 9–10**). These results support the reliability of the transcriptional patterns observed in the pathway array analyses.

**Figure 9.**
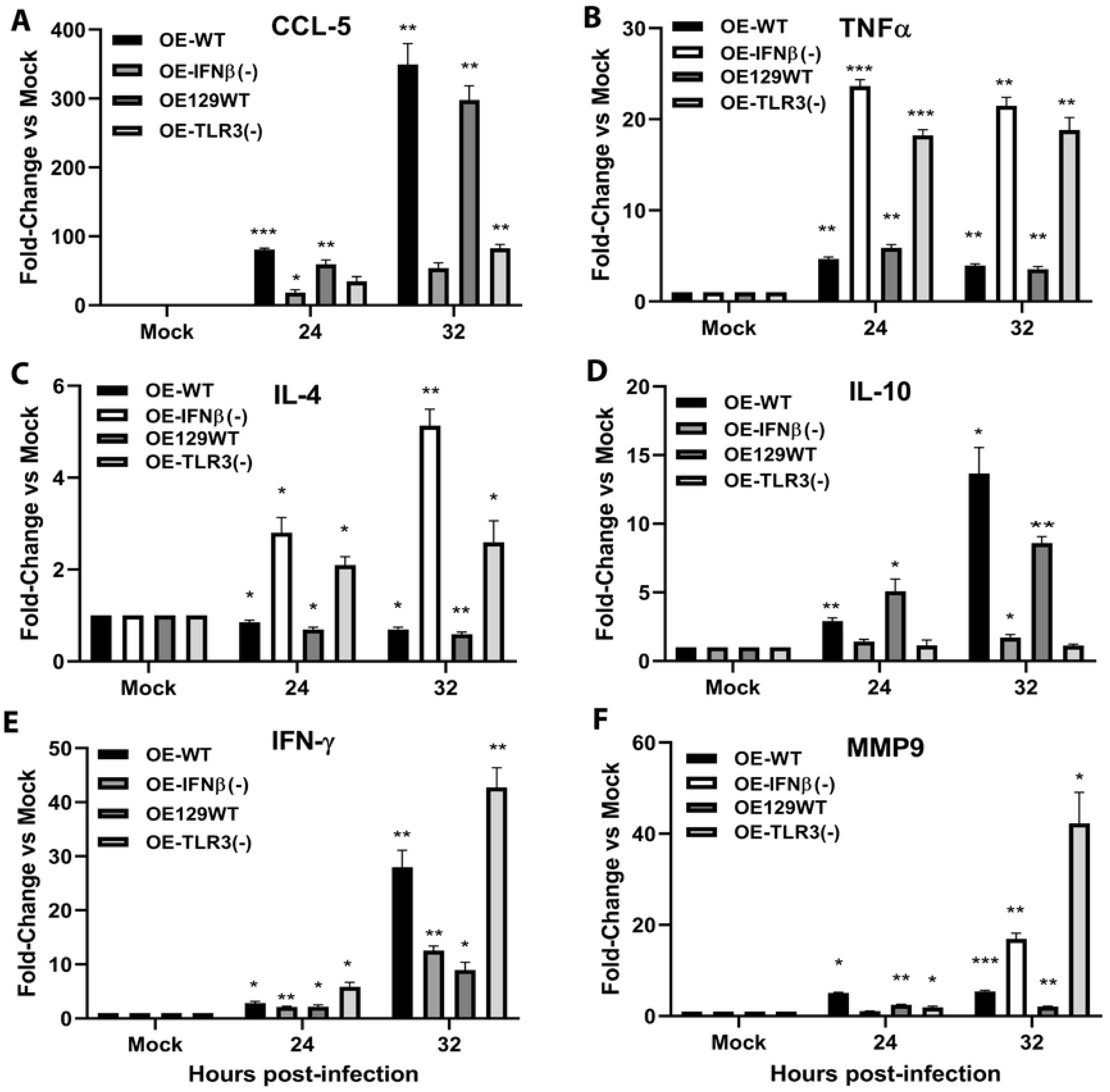
qPCR validation of selected immune response mediators involved in the innate and adaptive immune response, fibrosis, and scarring. Gene transcription levels of (A) CCL-5, (B) TNFα, (C) IL-4, (D) IL-10, (E) IFN-γ, and (F) MMP9 were measured by real-time quantitative PCR (qPCR) in the various wild-type and gene-deficient OE cells that were either mock-treated or infected with 10 IFU/cell *Chlamydia muridarum* for the time indicated. Data are representative of 3 independent experiments. *\*= p <0.05; **= p <0.005; ***= p <0.001 compared to mock control*.

**Figure 10.**
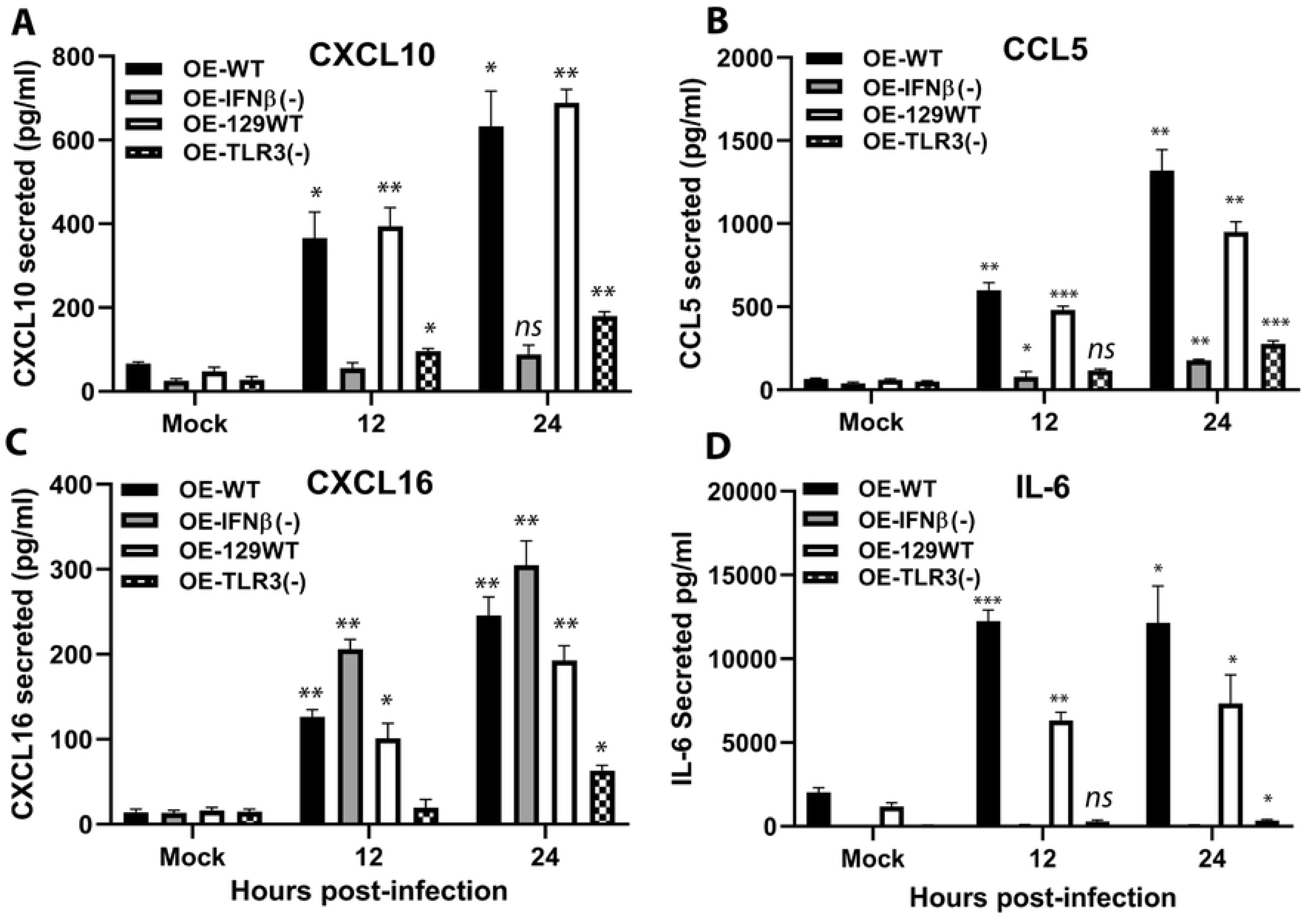
Independent qPCR validation of inflammatory pathway genes in wild-type versus IFNβ-deficient OE cells. ELISA was used to measure *Chlamydia-*infection-induced (A) CXCL10, (B) CCL5, (C) CXCL16, and (D) IL-6 secreted into the supernatants of wild-type and mutant OE cells that were either mock-treated or infected with the 10 IFU/ cell *C. muridarum*. Supernatants were collected at the indicated time post-infection. *\*\*\*= p<0.005; **= p<0.01; *= p<0.05; ns= not statistically significant when compared to mock-infected control cells.* The data presented are representative.

### Integrated model of IFN-β-dependent epithelial immune networks

Taken together, the transcriptional analyses described above support an integrated model in which IFN-β functions as a central regulatory hub coordinating epithelial immune responses during chlamydial infection (**Figure 11**). In this model, IFN-β signaling amplifies interferon-stimulated gene expression, promotes chemokine-mediated immune cell recruitment, and modulates inflammatory cytokine networks that influence tissue remodeling and fibrotic responses during infection.

**Figure 11.**
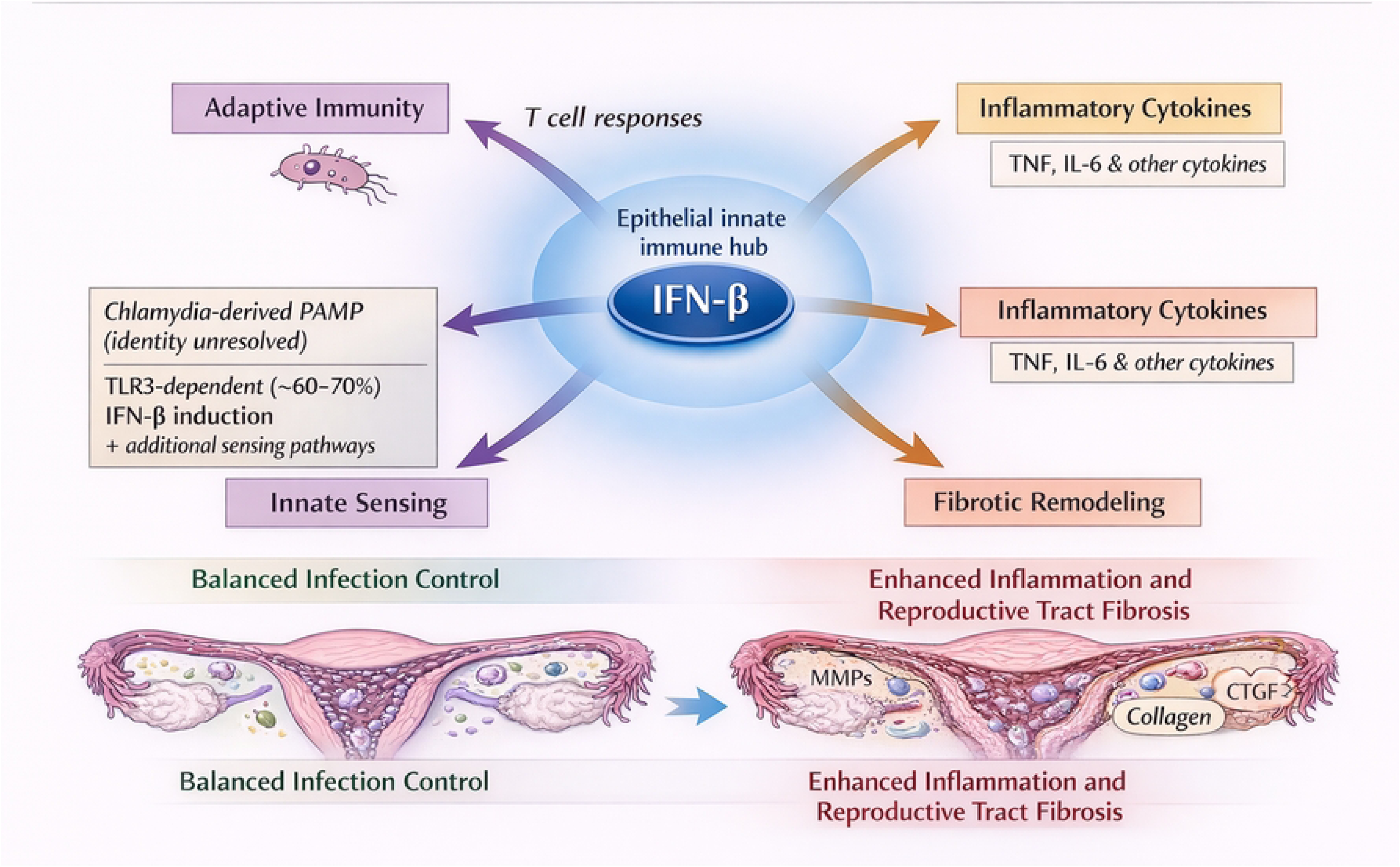
IFN-β orchestrates epithelial immune and fibrotic response networks during *Chlamydia muridarum* infection. Working model illustrating how epithelial cell–derived IFN-β integrates innate sensing pathways, inflammatory cytokine production, adaptive immune responses, and fibrotic remodeling during *Chlamydia muridarum* infection of the murine female reproductive tract. *Chlamydia*-induced IFN-β functions as a central epithelial immune hub downstream of TLR3-dependent and additional sensing pathways. Coordinated IFN-β signaling promotes balanced infection control and limits tissue pathology, whereas dysregulation of these pathways in the absence of IFN-β enhances inflammatory signaling and fibrotic remodeling, thereby increasing fibrosis in the reproductive tract.

## DISCUSSION

Collectively, these findings identify IFN-β as a central regulator of epithelial immune networks during chlamydial infection and suggest that disruption of IFN-β signaling may promote inflammatory and fibrotic pathology within the female reproductive tract. By coordinating transcriptional responses across multiple biological pathways---including innate immunity, interferon signaling, inflammatory cytokine networks, and fibrosis-associated pathways---IFN-β plays an important role in shaping epithelial cell immune responses to infection.

One of the most striking findings of this study is the extent to which IFN-β deficiency disrupts transcriptional responses across multiple immune pathways. In wild-type epithelial cells, infection induced coordinated activation of interferon-stimulated genes and chemokines involved in immune cell recruitment. In contrast, IFN-β deficiency resulted in widespread dysregulation of these transcriptional networks, including reduced induction of key interferon-responsive chemokines such as CCL5 and CXCL10. These findings are consistent with previous studies demonstrating that type-I interferons play important roles in coordinating host responses to intracellular pathogens (6). Consistent with the broader literature on chlamydial immunopathogenesis, these observations highlight the context-dependent roles of type-I interferon signaling, which can exert either protective or pathogenic effects depending on the cellular environment and stage of infection. In oviduct epithelial cells infected with Chlamydia, IFN-β appears to function as a transcriptional amplifier, enhancing the expression of interferon-stimulated genes involved in antimicrobial defense. Interestingly, IFN-β deficiency did not uniformly suppress inflammatory gene expression. While several interferon-associated chemokines were reduced in IFN-β-deficient cells, other inflammatory mediators---including TNF---were elevated. These findings suggest that IFN-β may function not only as an amplifier of epithelial immune responses but also as a regulatory signal that helps restrain excessive inflammatory signaling during infection.

The differential regulation of TNF and IL-6 observed in the present study highlights the complexity of inflammatory signaling networks during infection. Both cytokines have been implicated in inflammatory pathology and fibrosis in multiple disease contexts. (18, 34) However, the opposing patterns observed here suggest that IFN-β signaling may differentially regulate specific inflammatory pathways during infection.

One of the most notable findings of the present study is the impact of IFN-β deficiency on transcriptional programs associated with tissue remodeling and fibrosis. Several fibrosis-associated genes exhibited altered expression patterns in IFNβ-deficient cells, including genes involved in extracellular matrix regulation and pro-fibrotic signaling pathways. These observations are particularly relevant in the context of chlamydial pathogenesis. Chronic inflammation during genital chlamydial infection is strongly associated with the development of fibrotic lesions and scarring within the oviduct, processes that can ultimately impair reproductive function (2, 35-37).

Fibrosis usually results from dysregulated wound-healing responses in which inflammatory signaling promotes fibroblast activation and extracellular matrix protein deposition. Cytokines such as TNF, IL-6, and transforming growth factor (TGF)-associated pathways can influence these processes by regulating fibroblast activation and extracellular matrix turnover (38-41). Our transcriptional analyses revealed that IFN-β deficiency alters the expression of several genes associated with extracellular matrix remodeling, including *Ctgf (Ccn2), Tnf, Eng,* and *Serpine1*. CTGF (connective tissue growth factor) is a well-recognized mediator of fibrotic signaling that functions downstream of TGF-β signaling pathways to promote fibroblast activation and extracellular matrix deposition. CTGF has been implicated in the development of fibrosis in multiple organ systems (21, 42). Increased CTGF expression in IFNβ-deficient epithelial cells suggests that loss of IFN-β signaling may promote transcriptional programs associated with fibrotic tissue remodeling. In addition to CTGF-associated pathways, several matrix metalloproteinases (MMPs) were also dysregulated in IFNβ-deficient epithelial cells. MMPs regulate extracellular matrix turnover and tissue remodeling during inflammatory responses, and their dysregulation has also been implicated in fibrotic disease progression and tissue damage during chronic infection.

In conjunction with dysregulation of MMP and CTGF-associated signaling pathways, dysregulation of transcription pathways that affect TNF-α synthesis has also been widely implicated in tissue remodeling and fibrosis across multiple organ systems. TNF-α can promote inflammatory cascades that stimulate fibroblast activation and extracellular matrix production and is frequently elevated in chronic inflammatory diseases characterized by fibrotic pathology. Experimental studies have demonstrated that TNF signaling contributes to fibrosis in several disease models, including pulmonary, intestinal, hepatic, and renal fibrosis (43, 44). Elevated TNF-α signaling can activate downstream pathways that promote myofibroblast differentiation and extracellular matrix deposition during chronic inflammation. Consistent with these observations, our data show that TNF expression is increased in IFNβ-deficient epithelial cells during chlamydial infection, suggesting that IFN-β may normally function to restrain inflammatory circuits that promote fibrotic remodeling. Studies in multiple experimental systems have demonstrated that genetic or pharmacologic inhibition of TNF signaling can reduce inflammatory fibrosis, highlighting the central role of this cytokine in linking inflammation with tissue remodeling responses.

Interestingly, the inflammatory cytokine IL-6 exhibited a different pattern of regulation in our dataset, showing reduced expression in IFNβ-deficient cells despite increased TNF expression. TNF, IL-6, and other inflammatory mediators form interconnected cytokine networks that regulate fibroblast activation and extracellular matrix production during chronic inflammation (45). These cytokine networks function cooperatively to amplify inflammatory responses and influence fibrogenic signaling pathways. The divergent regulation of TNF and IL-6 observed in this study therefore highlights the complexity of cytokine signaling networks during infection. Rather than simply amplifying inflammation, IFN-β appears to regulate the balance among multiple inflammatory pathways, shaping the transcriptional environment in ways that may influence tissue repair versus fibrotic remodeling. Similarly, endoglin (*Eng*) is a co-receptor involved in TGF-β signaling pathways and plays important roles in vascular remodeling and fibrosis (46). Increased *Eng* expression observed in IFNβ-deficient cells may also reflect activation of signaling pathways that contribute to tissue remodeling responses during infection.

The altered regulation of *Serpine1* observed in this study is also noteworthy. SERPINE1 encodes plasminogen activator inhibitor-1 (PAI-1), a key regulator of fibrinolysis that influences extracellular matrix turnover and tissue remodeling (20). The marked reduction of *Serpine1* expression observed in infected OE-WT cells suggests that epithelial cells may actively regulate fibrinolytic pathways during *Chlamydia* infection. In contrast, persistence of *Serpine1* expression in IFNβ-deficient cells may alter the balance between matrix degradation and matrix deposition, potentially contributing to and exacerbating fibrotic remodeling processes. Taken together, these findings suggest that IFN-β signaling helps regulate the balance between inflammatory signaling and tissue remodeling pathways during infection. Dysregulation of these pathways in the absence of IFN-β may contribute to transcriptional programs associated with enhanced fibrotic responses.

Importantly, the presence of reproductive tract pathology in wild-type hosts indicates that chlamydial infection inherently triggers inflammatory responses capable of damaging host tissues. However, disruption of regulatory pathways such as IFN-β signaling may exacerbate pathological outcomes by altering the balance between protective immune responses and inflammatory processes that contribute to tissue remodeling and fibrosis. It is also important to recognize that IFN-β induction during chlamydial infection is not mediated exclusively by TLR3 signaling. Previous studies have shown that TLR3 deficiency reduces IFN-β production by approximately 60–70% *in vitro*, indicating that additional pathogen-sensing pathways contribute to interferon induction during infection (8, 9). These alternative pathways may include cytosolic nucleic acid sensors and other pattern recognition receptors that detect bacterial components during infection (32, 47-49).

Although the pathway-focused PCR arrays used in this study provide valuable insights into transcriptional responses during infection, several limitations should be considered. First, the arrays examine predefined gene sets and therefore do not capture the full complexity of the host transcriptional response. Second, the experiments were performed using non-transformed, but immortalized epithelial cell lines, which cannot fully recapitulate the complexity of the *in vivo* tissue environment. Importantly, these OE cell lines were derived directly from murine oviduct epithelial cells and retain key epithelial characteristics and innate immune signaling responses previously shown to recapitulate many aspects of epithelial responses observed during genital tract infection. Finally, the pathway-focused PCR arrays were selected for these studies because they enable targeted analysis of well-defined biological pathways while providing high sensitivity for detecting transcriptional changes in low-abundance immune mediators. This approach allowed focused examination of immune, interferon, inflammatory, and fibrosis-associated transcriptional programs most relevant to chlamydial pathogenesis. Nevertheless, we demonstrated consistency of transcriptional patterns observed across multiple pathway arrays, and our independent qPCR validation supports the robustness of the findings reported here. The independent qPCR analyses confirmed both the direction and magnitude of transcriptional changes observed in the pathway-focused PCR array datasets.

The collective findings of this study support a model in which IFN-β functions as a central regulatory node coordinating epithelial immune responses during chlamydial infection (**Figure 11**). In this model, infection of OE cells by *Chlamydia muridarum* activates multiple innate-immune sensing pathways that trigger IFN-β production. Our previous work has demonstrated that Toll-like receptor 3 (TLR3) signaling contributes substantially to IFN-β induction during infection; however, TLR3 deficiency reduces IFN-β production by approximately 60–70%, indicating that additional pathogen-sensing pathways also participate in interferon induction (9, 12). These observations suggest that epithelial IFN-β production results from the integration of multiple innate sensing pathways activated during intracellular infection.

Once produced, IFN-β activates transcriptional programs that influence several interconnected biological pathways. These include induction of interferon-stimulated genes involved in antimicrobial defense, activation of chemokine networks that promote recruitment of immune effector cells, regulation of inflammatory cytokine signaling, and modulation of extracellular matrix remodeling pathways associated with tissue repair. The coordinated regulation of these pathways likely plays a critical role in determining whether host immune responses effectively control infection while limiting tissue damage.

However, in the absence of IFN-β signaling, these regulatory networks become dysregulated. Reduced induction of interferon-associated chemokines such as CCL5 and CXCL10 may impair the recruitment of immune effector cells to infected tissues, while increased expression of inflammatory mediators such as TNF may promote inflammatory signaling pathways associated with tissue remodeling. Concurrent alterations in fibrosis-associated genes, including CTGF and SERPINE1, further suggest that dysregulated interferon signaling may influence extracellular matrix turnover and fibrotic responses during infection.

Importantly, the model presented here does not imply that wild-type hosts are protected from tissue pathology during infection. Rather, it has been well-demonstrated that genital tract infection with *Chlamydia muridarum* is known to produce significant inflammatory damage even in wild-type mice. However, our findings suggest that IFN-β signaling contributes to maintaining a regulatory balance among epithelial immune pathways. Our data presents the possibility that disruption of this balance may amplify inflammatory signaling networks and enhance transcriptional programs associated with tissue remodeling and fibrosis. Taken together, these findings support a model in which IFN-β acts as a central integrator of epithelial immune signaling at the interface of host defense during chlamydial infection, coordinating antimicrobial defense mechanisms while simultaneously regulating inflammatory and tissue remodeling responses that influence disease outcome.

An important question emerging from these findings concerns the potential contributions of other type-I interferons to chlamydial immunopathogenesis. Although the present study focused specifically on IFN-β, type-I interferon responses consist of a family of cytokines that signal through a shared receptor but can exhibit distinct biological functions depending on cellular context (6). Previous studies have suggested that global disruption of type-I interferon signaling can alter disease outcomes during *Chlamydia* infection, highlighting the complex roles of interferon pathways in regulating host responses to intracellular bacteria (50).

Our data suggest that IFN-β plays an important regulatory role in coordinating epithelial immune networks during infection. However, it is possible that other type-I interferons contribute differently to host responses during chlamydial infection. As an example, Interferon-α subtypes have been shown to exhibit distinct regulatory functions in inflammatory and antiviral signaling pathways. Differential regulations of IFNβ-dependent and IFNα-dependent signaling pathways may therefore influence the balance between protective immunity and inflammatory pathology during infection. Further investigation of how individual type-I interferons contribute to epithelial immune responses will be important for increasing our understanding the mechanisms underlying chlamydial immunopathogenesis. Dissecting the specific roles of IFN-β and IFN-α signaling pathways may ultimately provide new insight into how host immune responses influence reproductive tract pathology following infection.

In conclusion, the present study demonstrates that IFN-β functions as a central regulator of epithelial immune responses during chlamydial infection. By coordinating interferon signaling, inflammatory cytokine networks, and tissue-remodeling pathways, IFN-β helps maintain balanced host responses to infection. Disruption of these regulatory networks results in widespread transcriptional dysregulation that may promote inflammatory and fibrotic pathology within the female reproductive tract. These findings highlight IFN-β as a critical integrator of epithelial immune signaling during intracellular bacterial infection.

## Acknowledgements

This project was supported by National Institutes of Health Grants 1R01AI104944-01 and 1R21AI175847, and Marian University-Wood College of Osteopathic Medicine’s Office of Sponsored Programs and Research FRD grant funds. We graciously thank Michael S. Diamond at Washington University School of Medicine, Saint Louis, Missouri, USA for providing us with the use of the IFN-β KO mice we used to derive the OE cells used for these studies. We thank James Williams and staff at the Indiana University infectious disease diagnostic laboratory for assistance with the pathway-focused PCR arrays.

## Footnotes

1 This work was supported by National Institutes of Health Grants 1R01AI104944-01 and 1R21AI175847 (to W.A.D.).

## REFERENCES

1. Rowley J, Vander Hoorn S, Korenromp E, Low N, Unemo M, Abu-Raddad LJ, Chico RM, Smolak A, Newman L, Gottlieb S, Thwin SS, Broutet N, Taylor MM. 2019. Chlamydia, gonorrhoea, trichomoniasis and syphilis: global prevalence and incidence estimates, 2016. Bull World Health Organ 97:548–562P.

2. Brunham RC, Rey-Ladino J. 2005. Immunology of Chlamydia infection: implications for a Chlamydia trachomatis vaccine. Nat Rev Immunol 5:149–61.

3. Darville T, Hiltke TJ. 2010. Pathogenesis of genital tract disease due to Chlamydia trachomatis. J Infect Dis 201 Suppl 2:S114–25.

4. Derbigny WA, Hong SC, Kerr MS, Temkit M, Johnson RM. 2007. Chlamydia muridarum infection elicits a beta interferon response in murine oviduct epithelial cells dependent on interferon regulatory factor 3 and TRIF. Infect Immun 75:1280–90.

5. Johnson RM. 2004. Murine oviduct epithelial cell cytokine responses to Chlamydia muridarum infection include interleukin-12-p70 secretion. Infect Immun 72:3951–60.

6. McNab F, Mayer-Barber K, Sher A, Wack A, O’Garra A. 2015. Type I interferons in infectious disease. Nature Reviews Immunology 15:87.

7. Carrasco SE, et al. 2018. Toll-like receptor 3 (TLR3) promotes the resolution of Chlamydia muridarum genital tract infection in congenic C57BL/6N mice. PLoS One 13:e0195165.

8. Derbigny WA, et al. 2012. Identifying a role for Toll-like receptor 3 in the innate immune response to Chlamydia muridarum infection in murine oviduct epithelial cells. Infect Immun.

9. Derbigny WA, Johnson RM, Toomey KS, Ofner S, Jayarapu K. 2010. The Chlamydia muridarum-induced IFN-beta response is TLR3-dependent in murine oviduct epithelial cells. J Immunol 185:6689–97.

10. Kumar R, Derbigny WA. 2019. TLR3 Deficiency Leads to a Dysregulation in the Global Gene-Expression Profile in Murine Oviduct Epithelial Cells Infected with Chlamydia muridarum Madridge Journal of Bioinformatics and System Biology 1:1–13.

11. Kumar R, Gong H, Liu L, Ramos-Solis N, Seye CI, Derbigny WA. 2019. TLR3 deficiency exacerbates the loss of epithelial barrier function during genital tract Chlamydia muridarum infection. PLoS One 14:e0207422.

12. Derbigny WA, Shobe LR, Kamran JC, Toomey KS, Ofner S. 2012. Identifying a role for Toll-like receptor 3 in the innate immune response to Chlamydia muridarum infection in murine oviduct epithelial cells. Infect Immun 80:254–65.

13. Takaoka A, Mitani Y, Suemori H, Sato M, Yokochi T, Noguchi S, Tanaka N, Taniguchi T. 2000. Cross talk between interferon-gamma and -alpha/beta signaling components in caveolar membrane domains. Science 288:2357–60.

14. Hosey KL, Hu S, Derbigny WA. 2015. Role of STAT1 in Chlamydia-Induced Type-1 Interferon Production in Oviduct Epithelial Cells. J Interferon Cytokine Res 35:901–16.

15. Schachter J, Caldwell HD. 1980. Chlamydiae. Annu Rev Microbiol 34:285–309.

16. Schneider WM, Chevillotte MD, Rice CM. 2014. Interferon-stimulated genes: a complex web of host defenses. Annu Rev Immunol 32:513–45.

17. Attallah AM, Omran D, Omran MM, Albannan MS, Zayed RA, Saif S, Farid A, Hassany M, Yosry A. 2018. Fibro-Mark: a panel of laboratory parameters for predicting significant fibrosis in chronic hepatitis C patients. Br J Biomed Sci 75:19–23.

18. Igietseme JU, Omosun Y, Stuchlik O, Reed MS, Partin J, He Q, Joseph K, Ellerson D, Bollweg B, George Z, Eko FO, Bandea C, Liu H, Yang G, Shieh WJ, Pohl J, Karem K, Black CM. 2015. Role of Epithelial-Mesenchyme Transition in Chlamydia Pathogenesis. PLoS One 10:e0145198.

19. Karayi AK, Basavaraj V, Narahari SR, Aggithaya MG, Ryan TJ, Pilankatta R. 2020. Human skin fibrosis: up-regulation of collagen type III gene transcription in the fibrotic skin nodules of lower limb lymphoedema. Trop Med Int Health 25:319–327.

20. Sakai K, Matsuo J, Watanabe T, Okubo T, Nakamura S, Yamaguchi H. 2018. Subtle changes in host cell density cause a serious error in monitoring of the intracellular growth of Chlamydia trachomatis in a low-oxygen environment: Proposal for a standardized culture method. J Microbiol Methods 153:84–91.

21. Igietseme JU, Omosun Y, Nagy T, Stuchlik O, Reed MS, He Q, Partin J, Joseph K, Ellerson D, George Z, Goldstein J, Eko FO, Bandea C, Pohl J, Black CM. 2018. Molecular Pathogenesis of Chlamydia Disease Complications: Epithelial-Mesenchymal Transition and Fibrosis. Infect Immun 86.

22. Jin K, Ran J, Deng Y, Miao Z, Wei L, Yang Y, Yang M, Li T, Niu H, Yin G, Xie Q. 2025. Tumor Necrosis Factor-Induced Neuropilin-2 in Fibroblast-Like Synoviocytes Exacerbates Rheumatoid Arthritis. Arthritis Rheumatol doi:10.1002/art.70028.

23. Versteeg B, van den Broek LJ, Bruisten SM, Mullender M, de Vries HJC, Gibbs S. 2018. An Organotypic Reconstructed Human Urethra to Study Chlamydia trachomatis Infection. Tissue Eng Part A 24:1663–1671.

24. Xu Y, Zheng C, Jiang P, Ji S, Ullah S, Zhao Y, Su D, Xu G, Zhang M, Zou X. 2024. Fraxinellone alleviates colitis-related intestinal fibrosis by blocking the circuit between PD-1(+) Th17 cells and fibroblasts. Int Immunopharmacol 135:112298.

25. Lee HS, Kim WJ. 2022. The Role of Matrix Metalloproteinase in Inflammation with a Focus on Infectious Diseases. Int J Mol Sci 23.

26. Mustafa S, Koran S, AlOmair L. 2022. Insights Into the Role of Matrix Metalloproteinases in Cancer and its Various Therapeutic Aspects: A Review. Frontiers in Molecular Biosciences Volume 9–2022.

27. Bonnans C, Chou J, Werb Z. 2014. Remodelling the extracellular matrix in development and disease. Nature Reviews Molecular Cell Biology 15:786–801.

28. Lacy HM, Bowlin AK, Hennings L, Scurlock AM, Nagarajan UM, Rank RG. 2011. Essential role for neutrophils in pathogenesis and adaptive immunity in Chlamydia caviae ocular infections. Infect Immun 79:1889–97.

29. Belay T, Eko FO, Ananaba GA, Bowers S, Moore T, Lyn D, Igietseme JU. 2002. Chemokine and chemokine receptor dynamics during genital chlamydial infection. Infect Immun 70:844–50.

30. Poston TB, Lee DE, Darville T, Zhong W, Dong L, O’Connell CM, Wiesenfeld HC, Hillier SL, Sempowski GD, Zheng X. 2019. Cervical Cytokines Associated With Chlamydia trachomatis Susceptibility and Protection. J Infect Dis 220:330–339.

31. Shimada K, Porritt RA, Markman JL, O’Rourke JG, Wakita D, Noval Rivas M, Ogawa C, Kozhaya L, Martins GA, Unutmaz D, Baloh RH, Crother TR, Chen S, Arditi M. 2018. T-Cell-Intrinsic Receptor Interacting Protein 2 Regulates Pathogenic T Helper 17 Cell Differentiation. Immunity 49:873–885 e7.

32. Nagarajan UM, Sikes J, Prantner D, Andrews CW, Jr., Frazer L, Goodwin A, Snowden JN, Darville T. 2011. MyD88 deficiency leads to decreased NK cell gamma interferon production and T cell recruitment during Chlamydia muridarum genital tract infection, but a predominant Th1 response and enhanced monocytic inflammation are associated with infection resolution. Infection and Immunity 79:486–98.

33. García-Domínguez M. 2025. The Role of IL-23 in the Development of Inflammatory Diseases. Biology. 14(4):347. doi:10.3390/biology14040347.

34. Zhao X, Zhao Q, Zhu X, Huang H, Wan X, Guo R, Zhao Y, Chen D, Xu D. 2020. Study on the correlation among dysbacteriosis, imbalance of cytokine and the formation of intrauterine adhesion. Ann Transl Med 8:52.

35. Jiang J, Karimi O, Ouburg S, Champion CI, Khurana A, Liu G, Freed A, Pleijster J, Rozengurt N, Land JA, Surcel HM, Tiitinen A, Paavonen J, Kronenberg M, Morre SA, Kelly KA. 2012. Interruption of CXCL13-CXCR5 axis increases upper genital tract pathology and activation of NKT cells following chlamydial genital infection. PLoS One 7:e47487.

36. Rank RG, Sanders MM, Kidd AT. 1993. Influence of the estrous cycle on the development of upper genital tract pathology as a result of chlamydial infection in the guinea pig model of pelvic inflammatory disease. Am J Pathol 142:1291–6.

37. Wyrick PB. 2010. Chlamydia trachomatis persistence in vitro: an overview. J Infect Dis 201 Suppl 2:S88–95.

38. Smith LC, Moreno S, Robertson L, Robinson S, Gant K, Bryant AJ, Sabo-Attwood T. 2018. Transforming growth factor beta1 targets estrogen receptor signaling in bronchial epithelial cells. Respir Res 19:160.

39. Miller CG, Budoff G, Prenner JL, Schwarzbauer JE. 2017. Minireview: Fibronectin in retinal disease. Exp Biol Med (Maywood) 242:1–7.

40. Tschumperlin DJ. 2015. Matrix, mesenchyme, and mechanotransduction. Ann Am Thorac Soc 12 Suppl 1:S24–9.

41. Hubmacher D, Apte SS. 2013. The biology of the extracellular matrix: novel insights. Curr Opin Rheumatol 25:65–70.

42. Burton MJ, Rajak SN, Hu VH, Ramadhani A, Habtamu E, Massae P, Tadesse Z, Callahan K, Emerson PM, Khaw PT, Jeffries D, Mabey DC, Bailey RL, Weiss HA, Holland MJ. 2015. Pathogenesis of progressive scarring trachoma in Ethiopia and Tanzania and its implications for disease control: two cohort studies. PLoS Negl Trop Dis 9:e0003763.

43. Steele H, Cheng J, Willicut A, Dell G, Breckenridge J, Culberson E, Ghastine A, Tardif V, Herro R. 2023. TNF superfamily control of tissue remodeling and fibrosis. Frontiers in Immunology Volume 14–2023.

44. Wynn TA, Ramalingam TR. 2012. Mechanisms of fibrosis: therapeutic translation for fibrotic disease. Nat Med 18:1028–40.

45. Alcolea MPI, Starita Fajardo G, Pena Rodriguez M, Lucena Lopez D, Suarez Carantona C, Lopez Paraja M, Garcia de Vicente A, Viteri-Noel A, Gonzalez Garcia A. 2025. Advances in the Molecular Mechanisms of Pulmonary Fibrosis in Systemic Sclerosis: A Comprehensive Review. Int J Mol Sci 26.

46. Huang Q, Xiao R, Lu J, Zhang Y, Xu L, Gao J, Sun J, Wang H. 2022. Endoglin aggravates peritoneal fibrosis by regulating the activation of TGF-beta/ALK/Smads signaling. Front Pharmacol 13:973182.

47. Archer KA, Durack J, Portnoy DA. 2014. STING-dependent type I IFN production inhibits cell-mediated immunity to Listeria monocytogenes. PLoS Pathog 10:e1003861.

48. Prantner D, Darville T, Nagarajan UM. 2010. Stimulator of IFN gene is critical for induction of IFN-beta during Chlamydia muridarum infection. J Immunol 184:2551–60.

49. Tanaka Y, Chen ZJ. 2012. STING specifies IRF3 phosphorylation by TBK1 in the cytosolic DNA signaling pathway. Sci Signal 5:ra20.

50. Nagarajan UM, Prantner D, Sikes JD, Andrews CW, Jr., Goodwin AM, Nagarajan S, Darville T. 2008. Type I interferon signaling exacerbates Chlamydia muridarum genital infection in a murine model. Infect Immun 76:4642–8.

